# Species-specific genetic routes to a shared phenotype: convergent adaptation of beans to European long days

**DOI:** 10.1101/2025.09.14.676158

**Authors:** M Rendón-Anaya, L Yu, Xiao-Feng Yang, PK Ingvarsson

## Abstract

The loss of photoperiod sensitivity was a key prerequisite for the adaptation of common bean (*Phaseolus vulgaris*) and scarlet runner bean (*P. coccineus*) to European environments. While the genetic architecture of this trait has been studied for decades, primarily through QTL mapping in RILs and NILs, its molecular basis remains incompletely resolved. Here, we combine whole-genome resequencing of over 200 *P. vulgaris* accessions spanning Mesoamerican, Andean, and European origins with 130 *P. coccineus* cultivars from Mesoamerica and Europe. Phenotyping under controlled long-day conditions revealed distinct distributions of photoperiod sensitivity across species and gene pools. Genome-wide association and haplotype analyses identified single nucleotide variants strongly linked to flowering under long days. Consistent with prior studies, signals of selection reflect independent domestication events in the Americas. Importantly, we detect species-specific selection on core photoperiod regulators, demonstrating that both species achieved similar adaptive outcomes via functionally convergent genetic routes. By explicitly accounting for major loci shaping growth habit in our analyses, we were able to unmask additional genomic regions contributing to flowering responses in European cultivars. Together, these results show that adaptation to long-day environments occurred through stepwise, species-specific genomic innovations, first during domestication in the Americas and later under European cultivation. Our findings highlight the role of pathway-level convergent evolution in shaping complex domestication traits and underscore the potential of parallel genomic strategies for crop adaptation.

## Introduction

Many aspects of plant growth and development show fine-tuned local adaptation, and this is particularly true for the transition from vegetative growth to flowering (Roux, et al. 2006). In many plant species, flowering requires exposure to specific photoperiods and/or temperatures, and reproduction may be delayed or prevented when these requirements are not met. As plants colonize new environments, genetic changes that modify or relax these constraints can enable range expansion across diverse climatic regimes. Although flowering time can be considered a relatively simple developmental switch, the transition from vegetative to reproductive growth entails coordinated changes in stem elongation, apical dominance, lateral branching, and resource allocation. Numerous genes contributing to the control of flowering have been identified in *Arabidopsis* and other model species (Weller and Ortega 2015; Blackman 2017), and the underlying regulatory networks are broadly conserved across angiosperms (Wu, et al. 2024).

The genus *Phaseolus* provides a unique framework to investigate the genetic basis of flowering adaptation. Common bean (*P. vulgaris*), a predominantly selfing annual species, originated in Mesoamerica and comprises two major gene pools: Mesoamerican and Andean, which diverged ∼165,000 years ago (Schmutz, et al. 2014). Independent domestications occurred in both regions ∼8,000 years ago, producing parallel domestication syndromes. Following introduction to Europe after the Columbian exchange, *P. vulgaris* rapidly adapted to higher latitudes and cooler temperatures, with European cultivars showing predominantly Andean ancestry (Angioi, et al. 2010b). Populations segregate for strong short-day responses, day-neutral flowering, or long-day tolerance, and European landraces clearly reflect selection for flowering under long-day conditions (>14 h), consistent with adaptation to northern latitudes. In common bean, several loci influencing photoperiod sensitivity have been mapped, including *TERMINAL FLOWER 1* (Gonzalez, et al. 2016), *PHYTOCHROME A* (Weller, et al. 2019), as well as CONSTANS-like and AGAMOUS-like 8 candidates (Gonzalez, et al. 2021), highlighting the polygenic nature of this trait.

Runner bean (*P. coccineus*), a perennial climbing species also domesticated in Mesoamerica, shares this trajectory of introduction and cultivation in Europe but differs markedly in its reproductive biology and demographic history. Adaptation of *P. coccineus* to European environments has been shaped by founder events and subsequent differentiation in flowering phenology along latitudinal gradients (Rodriguez, et al. 2013). Yet, unlike *P. vulgaris*, the genetic determinants of photoperiod sensitivity in *P. coccineus* remain largely unexplored. Insights from legumes more broadly implicate conserved photoperiod and circadian regulators, including *FT* homologues, *PHYA*, *ELF3*, and the legume-specific *E1* gene (Weller & Ortega 2015), suggesting that runner bean adaptation could involve both conserved pathways and novel, species-specific solutions.

Because the loss of photoperiod sensitivity has independently arisen in both the Mesoamerican and Andean gene pools of *P. vulgaris* and has been essential for the adaptation of *P. coccineus* in Europe, these two species together provide an exceptional system to test hypotheses of parallel versus convergent evolution at the molecular level. Convergent traits may arise through unrelated genes in distinct pathways, whereas parallelism reflects repeated changes in orthologous genes or regulatory modules (Pickersgill 2018; Woodhouse and Hufford 2019). Distinguishing between these evolutionary processes is central to understanding the genetic routes available to crops during adaptation.

Here, we integrate whole-genome sequencing of 232 *P. vulgaris* accessions of diverse origin with 130 *P. coccineus* cultivars from Mesoamerica and Europe. Using genome-wide scans of selection, GWAS, and haplotype structure analyses, we dissect the genomic basis of photoperiod sensitivity in both species. This comparative framework allows us to identify gene-pool-specific signatures of adaptation in common bean, uncover previously uncharacterized determinants in runner bean, and evaluate the extent to which the repeated loss of photoperiod sensitivity reflects parallel versus convergent evolution during Phaseolus domestication and adaptation to Europe.

## Materials and methods

### Plant material and growing conditions

We collected a diverse panel of 232 *Phaseolus vulgaris* accessions of Mesoamerican, Andean and European origin, that include commercial accessions, land races and wild-collected individuals from the centres of origin. We also built a panel of 130 *Phaseolus coccineus* accessions of Mesoamerican and European origins. These accessions are publicly available at the International Centre for Tropical Agriculture in Cali, Colombia; the Nordic Genetic Resource Center and the European Search Catalogue for Plant Genetic Resources at Gatersleben, Germany.

We phenotyped our panel of *Phaseolus* accessions in response to daylength as follows: the accessions were sown in medium pots (10 cm of diameter) with 750 gr of sterile soil. To avoid temperature as a confounding variable, plants were grown in phytotron chambers under the following regime: Short days: 10hrs light; 18℃ night/22 ℃ day. Long days, 16hrs light, 18℃ night/22 ℃ day. We counted the days from germination to the emergence of the first flower buds in both conditions. Those accessions that flowered under long and short day-lengths were labeled as photoperiod insensitive (PhIns), while those flowering only under short days, were labeled photoperiod sensitive (PhSens). This classification was then coded as a binary trait (0 for PhSens, 1 for PhIns) for the association analyses. We grew two plants per accession in each condition at the Plant Cultivation Facility at the BioCentre, Swedish University of Agricultural Sciences, Uppsala.

### Genome re-sequencing, mapping and SNP calling

Total genomic DNA was extracted from frozen leaf tissue for all individuals using the DNeasy plant mini prep kit (QIAGEN, Valencia, CA, USA). Briefly,1 µg of high-quality DNA was used for paired-end libraries construction. The libraries (TruSeq, PCR-free, 350bp) were subjected to paired-end sequencing (2x150) at the National Genomics Infrastructure at Science for Life Laboratory, Stockholm, on an Illumina NovaSeq 6000 to a mean per-sample depth of approximately 15X.

Raw reads were mapped to the reference genome of *P. vulgaris* (https://phytozome-next.jgi.doe.gov/info/Pvulgaris_v2_1) using BWA-mem version 2.2.3 (Li and Durbin 2009). Aligned reads were flagged for duplicates using the MarkDuplicates program in Picard tools version 1.119 (http://broadinstitute.github.io/picard/). Multi-sample single nucleotide polymorphism (SNP) calling was performed using the Genome Analysis Toolkit (GATK) version 3.8. For GATK, SNPs were called using the tools HaplotypeCaller and GenotypeGVCF. SNPs were only retained if they matched the following criteria: QualByDepth > 20 (a measure of alternative allele quality independent of read depth), MQ > 30 (the minimum phred-scaled mapping quality of the reads supporting the alternative allele), FisherStrand < 10 (whether an alternative allele was predominantly supported by one read orientation only), −2 < BaseQualityRankSumTest < 2 (a test statistic to assess whether the base quality of reads supporting the alternative allele was significantly worse than reads supporting the reference allele), −2 < ReadPosRankSumTest < 2 (a test statistic to assess whether the base position in a read supporting the alternative allele was significantly different than the base position in a read supporting the reference allele), −2 < MappingQualityRankSumTest < 2 (a test statistic to assess whether the mapping quality of reads supporting the alternative allele was significantly worse than reads supporting the reference allele), RMSMappingQuality > 30 (an estimation of the overall mapping quality of reads supporting an alternative allele).

We calculated genome-wide breadth and depth of coverage using samtools (Li, et al. 2009). After obtaining the average coverage in our collection, we set a minimum and maximum number of reads per site to filter out very low/high depth sites. For downstream analyses we kept only biallelic sites with no more that 30% missingness in the vcf file, which were obtained with BCFtools (Li 2011).

### Population genomics

In order to remove potentially duplicated samples between genebanks and related individuals we calculated the identity by descent (IBD) with PLINK v1.9 (Purcell, et al. 2007). Samples with a PI_HAT > 0.4 were removed. SNP-based PCAs were constructed using plink –pca on pruned, unlinked sites with a minimum MAF exceeding 0.05. Next, we used vcftools to output the genotype likelihood information contained in the pruned VCF file; the resulting beagle-formatted file was input into NGSAdmix (Skotte, et al. 2013) to estimate individual admixture proportions across a varying number of ancestral populations (K=2 to K=5). To calculate standard population genomics estimators between subpopulations (e.g. domesticated vs wild, or photoperiod sensitive vs insensitive) such as pi, F_ST_ and dxy we used the python package *genomics_general* defining 10Kb, non-overlapping windows (https://github.com/simonhmartin/genomics_general).

Linkage-disequilibrium (LD) decay was plotted with PopLDdecay (Zhang, et al. 2019). In a downstream analysis, haplotype structure surrounding significant GWAS SNPs was assessed using LDBlockShow v1.40 for both *Phaseolus vulgaris* and *P. coccineus*. Prior to LD calculation, SNP datasets were filtered to retain only variants with a minor allele frequency (MAF) ≥ 0.05 to reduce the influence of rare alleles. For each species, genomic regions spanning ±100 kb around the focal GWAS SNPs were extracted, and pairwise LD (r²) was calculated between all SNPs within each window. Haplotype blocks were defined based on the confidence interval method implemented in LDBlockShow, and LD heatmaps were generated to visualize the extent of local haplotype structure. Comparisons were made between photoperiod-insensitive and photoperiod-sensitive accessions to assess differences in LD patterns, considering the contrasting mating systems of the two species (selfing in *P. vulgaris* versus predominantly outcrossing in *P. coccineus*).

For *P. vulgaris*, we produced an ancestral genome based on the most represented variants in the wild Mesoamerican *P. vulgaris* samples using ANGSD -doFasta 2 that outputs a fasta file taking the most common base per site given a series of bam infiles mapped to the reference genome (Korneliussen, et al. 2014). With this, we could extrapolate the ancestral allele in the vcf files for further analyses.

### Alleles favored by evolution or positive selection

We scanned the genome of *P. coccineus* for signals of positive selection using “integrated selection of alleles favoured by evolution”, iSAFE (Akbari, et al. 2018). This method first calculates haplotype allele frequency scores based on the presence of derived alleles in a particular haplotype, which is then used to calculate SAFE scores. These SAFE scores are in turn calculated across a region of given size in 50% overlapping windows of 300 bp to culminate in an iSAFE signal. These statistics can be calculated for large regions of phased haplotypes, which we obtained with BEAGLE v.4.1 (Browning and Browning 2007), so chromosomes were divided into 3 Mb windows for each iSAFE iteration. The iSAFE software can be set to run under a case-control mode, with the case populations being the group of photoperiod insensitive cultivars, and the control population being the cultivars that can only flower under short days and are therefore, photoperiod sensitive. This method was not used on P*. vulgaris* due to the selfing nature.

### Genome wide associations and haplotype differentiation

We coded the phenotypic data as binary matrices, i.e. accessions classified as photoperiod insensitive were coded with 1, whereas those classified as sensitive were coded with a 0. We used SNP panels of pruned sites (filtered with plink -maf 0.05 -indep-pairwise 100 10 0.2 -geno 0.1) to run genome-wide association analyses. We converted each vcf file to a matrix of 0,1 and 2 values for homozygous (ref/alt) or heterozygous genotypes with vcftools (vcftools –012). As we allowed 10% of missingness in this genotype filtering step, we had to impute the missing genotype information to avoid numeric biases in the GWAS calculations. For each column in the matrix representing individual positions, we calculated the mean genotype value (not including missing genotypes coded as -1) that we used to fill the missing genotypes. This numeric matrix was the input for GWAS analyses.

We combined genotype and phenotypic data on photoperiod sensitivity through genome-wide association mapping (GWAS). GWAS was performed using several models: mixed linear model (MLM), multiple loci mixed model (MLMM), and Bayesian-information and linkage-disequilibrium iteratively nested keyway (BLINK), all implemented in GAPIT3 (Wang and Zhang 2021). We controlled for the effects of population structure by setting the number of relevant PCs at 5 for *P. vulgaris* and 4 for *P. coccineus*. Furthermore, we added the growth habit (determinate/indeterminate) and seed-bank origin as covariates in *P. vulgaris* and *coccineus*, respectively.

## Results

### Population structure

We re-sequenced 232 *P. vulgaris* accessions of Mesoamerican, Andean and European origin, that include commercial accessions, land races and wild-collected individuals from the centres of origin. The estimated average depth and breadth of coverage were 13,7X and 90.2% across the samples. Due to strong relatedness and to possible duplications of some accessions across seedbanks, we retained 172 accessions for downstream analyses.

Based on 161,391 pruned SNPs we obtained a PCA that grouped the accessions following their genetic background: PC1 separates the accessions in the two dominant gene pools, Mesoamerican (MA) and Andean (AN), while PC2 separates them according to their domesticated/wild state (Figure 1A). We observe two divergent groups of wild accessions, two clusters of domesticated accessions in proximity to their wild ancestors, which confirms both independent domestication events, and finally, a large group of accessions grown in Europe that can be separated according to their genomic background, but that also displays clear signs of admixture following the introduction of the species in Europe. It is clear that the majority of accessions grown in Europe had a preferential Andean background, which confirms previous observations regarding the gene pools present in the continent (Angioi, et al. 2010a).

**Figure 1.**
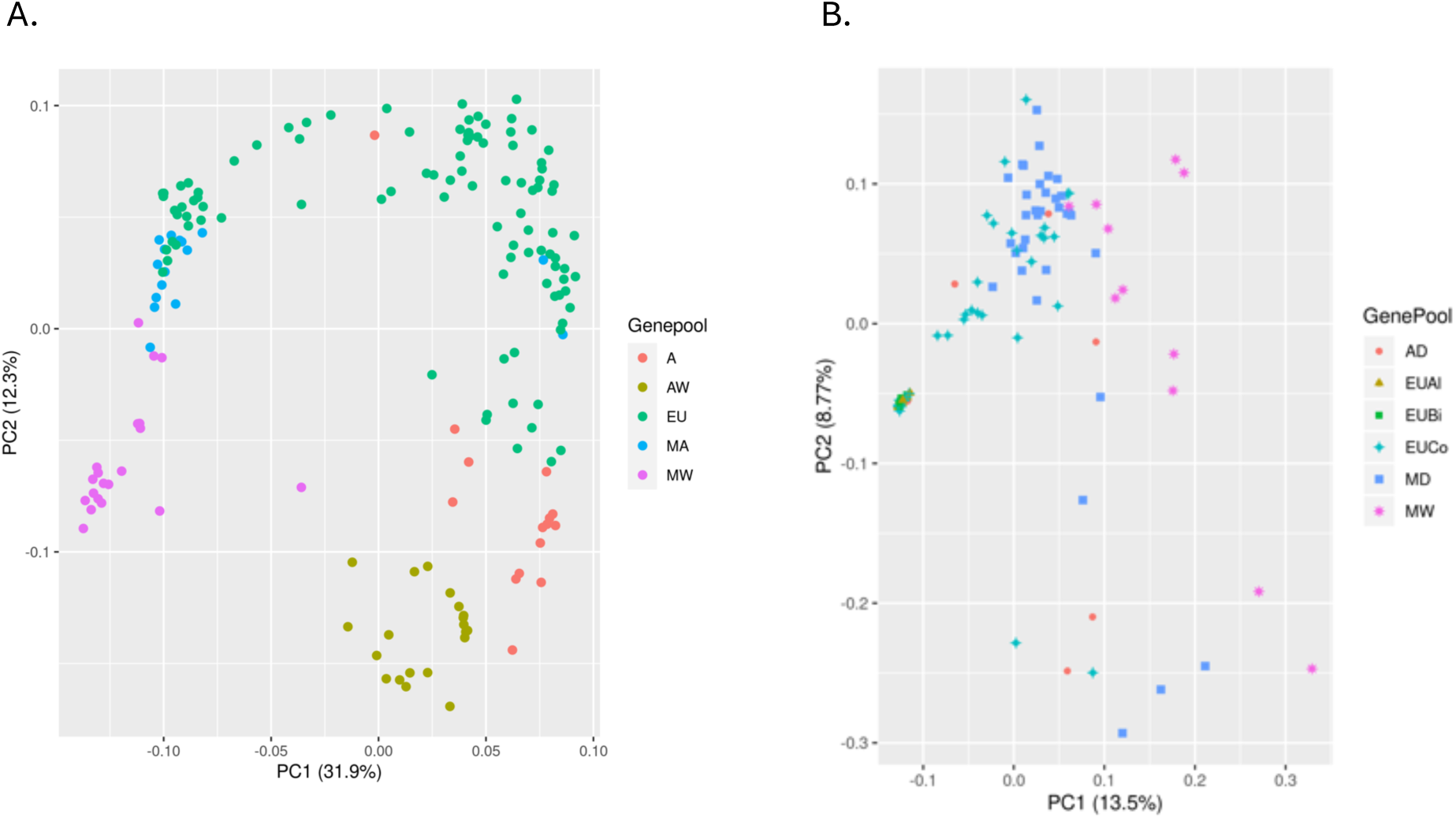
Population structure. SNP-based PCAs were constructed using non-rare, unlinked variants across the 11 chromosomes in *P. vulgaris* (A) *P. coccineus* (B).

Within our panel of runner bean accessions, the PCA based on 84122 pruned sites shows a less structured population from the Americas, but a very compact cluster of European cultivars (Figure 1B). Furthermore, this tight cluster corresponds to cultivars we obtained from IPK and does not include accessions collected in Europe stored at CIAT. To gain further resolution, we ran NGSAdmix and confirmed a very homogeneous group of *P. coccineus* cultivars from European seed banks, that shows no admixture with any of the American relatives. At a K=2, we clearly see the Mesoamercian origin of the European cultivars; however, as K increases, it is evident that the European-labelled accessions from CIAT show clear signs of admixture with other Mesoamerican domesticated cultivars (Suppl. Figure 1).

### Mating system differences

Our analyses confirmed the contrasting mating systems of the two species. *Phaseolus vulgaris*, a predominantly selfing species, displayed slow linkage disequilibrium (LD) decay, with significant correlations between markers extending over long physical distances. In contrast, *P. coccineus*, an obligate outcrosser, exhibited rapid LD decay consistent with its higher effective recombination rate. Estimates of the inbreeding coefficient further supported these differences, with *P. vulgaris* accessions showing high values indicative (>0.5) of selfing, while *P. coccineus* exhibited coefficients close to zero, consistent with extensive outcrossing. These results corroborate the expected reproductive biology of the two species and provide a genomic baseline for interpreting GWAS signals (Suppl. Figure 2).

#### Growth habit associated haplotypes in the common bean

We ran genome wide association analyses for determinacy in *P. vulgaris* (Figure 2) using both multi-locus models, MLMM and Blink, as well as single locus model, MLM, all implemented in GAPIT3; the advantages of these models in terms of statistical power vs computational cost have been discussed elsewhere (Wang and Zhang 2021). All methods detected a significant, dominant signal around 44.8Mb on chromosome 1 associated to differences in growth habit (Figure 3) that matches the previously identified *Fin* locus (Kwak, et al. 2008; Kwak, et al. 2012). At this locus, the highest scoring SNPs Chr01_44852374_G_A (Blink p-value of 9.22e-112, maf=0.39) and Chr01_45392359_C (p-value 1.78e-105, maf=0.08) explain 22.5 and 44.7% of the phenotypic variance, and are located in close proximity to *Terminal Flower 1* (*TFL1*, Phvul.001G189200 located between 44856139 and 44857862bp), which has previously shown to be responsible for the indeterminate phenotype in *Phaseolus vulgaris* (Koinange, et al. 1996).

**Figure 2.**
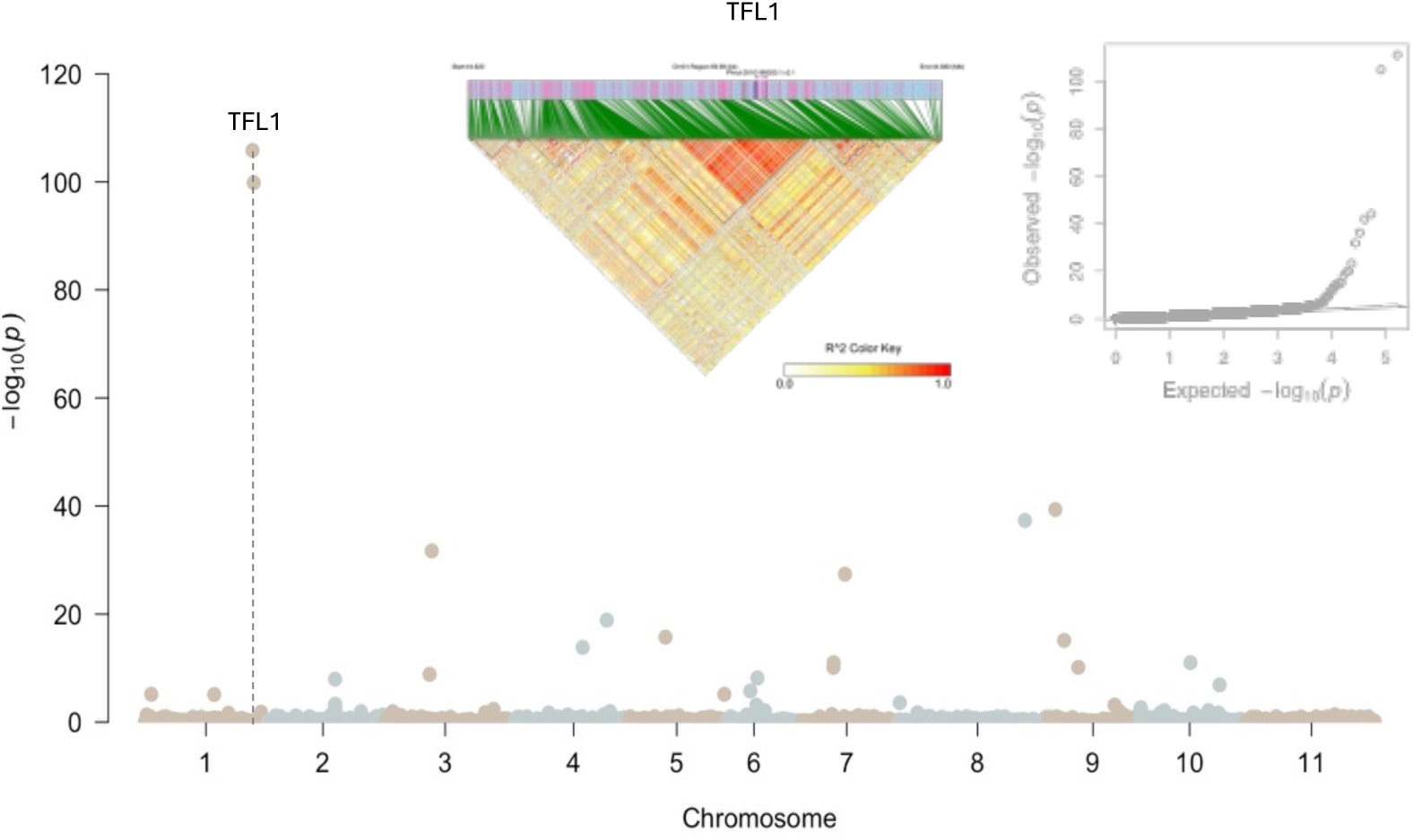
Genome-wide association analysis of determinacy in *Phaseolus vulgaris*. (A) Manhattan plot showing SNP associations across the 11 chromosomes. GWAS was performed using the BLINK model implemented in GAPIT3. The strongest association maps to the TFL1 (Fin) locus on chromosome 1, consistent with its known role in controlling determinacy and growth habit. Additional smaller peaks were observed elsewhere in the genome but did not colocalize with previously characterized determinacy genes. (B) Quantile–quantile (Q–Q) plot of observed versus expected *p*-values, showing good overall model fit and a pronounced excess of low *p*-values at the tail, consistent with the strong genetic effect of TFL1. (C) Linkage disequilibrium (LD) heatmap and haplotype block structure around the TFL1 locus.

**Figure 3.**
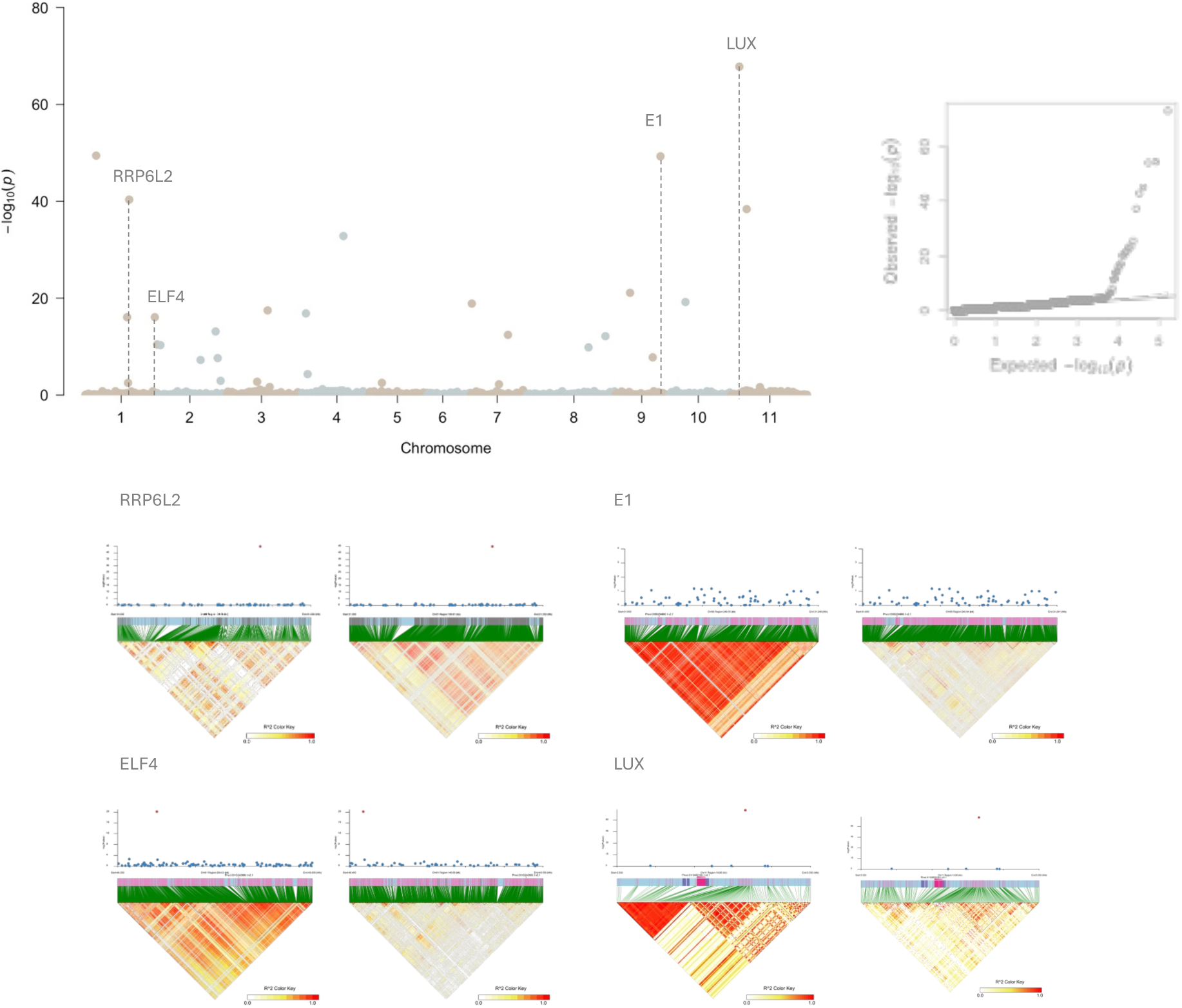
Genome-wide association analysis of photoperiod sensitivity in *Phaseolus vulgaris*. (A) Manhattan plot showing SNP associations across the 11 chromosomes. GWAS was performed using the BLINK model implemented in GAPIT3. Significant peaks were detected on chromosomes 1, 9, and 11. On chromosome 1, signals were located near ELF4, a circadian regulator; on chromosome 9, near E1, a repressor of *FT* orthologs under long days; and on chromosome 11, near LUX, a component of the evening complex. (B) Quantile–quantile (Q–Q) plot of observed versus expected *p*-values, demonstrating good model fit with enrichment of low *p*-values at the tail, consistent with true associations. (C) Linkage disequilibrium (LD) heatmaps and haplotype block structures for the main GWAS peaks. Extended haplotype blocks were observed around significant SNPs on chromosomes 1, 9, and 11, reflecting the selfing mating system of *P. vulgaris* and supporting the robustness of the associations. PhIns (left panel); PhSens (right panel).

#### GWAS on photoperiod sensitivity in the common bean

Because of the proximity between Fin and Ppd in chromosome 1, and due to the overlap between changes in growth habit and photoperiod responses along domestication, we used the binary encoded growth habit (determinate/indeterminate) as a covariate for the identification of new signals behind photoperiod response differences in *P. vulgaris*. Genome-wide association analysis using Blink revealed several significant SNP-trait associations distributed primarily across chromosomes 01, 09, and 11. On chromosome 01, significant signals were detected in close proximity to RRPL62 (Phvul.001G114400; SNP Chr01_31147588_C_T, p-value 1.14E-45) and ELF4 (Phvul.001G242900, SNP Chr01_49410758_T_C, p-value 6.35E-21). These sites explain 10.9 and 3.7% of the phenotypic variance, respectively. The *RRPL62* gene encodes a putative pentatricopeptide repeat-like protein, which may be involved in post-transcriptional regulation and has been associated with developmental timing in other plant systems. The *ELF4* (*EARLY FLOWERING 4*) gene is a well-characterized component of the circadian clock that regulates photoperiodic flowering by stabilizing evening-phased oscillations and interacting with other clock proteins to fine-tune *FT* expression (Wang, et al. 2024).

On chromosome 09, a strong association was identified near the E1 locus (Phvul.009G204600 ; SNP Chr09_31884646_A_G, p-value 9.72E-55), a major flowering repressor originally described in soybean (Fang, et al. 2024). This site is found at a high frequency (maf=0.37) and explains 6.2% of the phenotypic variance. The *E1* gene encodes a B3-type transcription factor that inhibits the expression of *FT* orthologs under long days, thereby delaying flowering; the presence of a homologous locus in *P. vulgaris* suggests conservation of photoperiod-regulatory pathways across legumes. Although the marker and the E1 locus are ∼800Kb apart, the LD pattern in photoperiod insensitive cultivars, indicates the presence of a strong haplotype block that spans the region.

On chromosome 11, a significant SNP was detected in close physical proximity to LUX ARRHYTHMO (Phvul.011G0621; SNP Chr11_5543230_T_A, p-value 1.01E-73), another circadian clock gene. This SNP was found at a frequency of maf=0.08, explaining 16.2% of the phenotypic variance. *LUX* is a central component of the evening complex, acting together with *ELF3* and *ELF4* to regulate circadian rhythms and flowering induction (Liew, et al. 2017). Its identification here underscores the importance of clock genes in shaping flowering time variation in *Phaseolus*. Additional signals of association were detected on other chromosomes; however, these did not colocalize with annotated flowering or photoperiod-related genes.

To verify the robustness of GWAS signals, linkage disequilibrium (LD) around the focal SNPs was analyzed using LDBlockShow. Strong LD blocks were observed around SNPs near RRPL62, ELF4, E1, and LUX in photoperiod-insensitive cultivars, indicating tight haplotype structure in these regions. In contrast, photoperiod-sensitive cultivars exhibited little to no LD at the same loci (Figure 3).

Finally, we detected some overlaps between GWAS signals and Fst outliers (top 5%; Suppl. Figure 3). For example, ELF4 on chromosome 1 is encoded within a highly differentiated region between wild and domesticated populations of the Mesoamerican and Andean gene pools, and between domesticated and European populations from the Mesoamerican but not the Andean gene-pool. Interestingly, the Fst values between Andean and Mesoamerican domesticated populations around this locus is as high as 0.58, suggesting strongly different haplotypes between gene-pools. Another interesting overlap occurs on chromosome 11, where the region spanning 5.3-5.6Mb that harbors LUX, is strongly differentiated as a result of the adaptation to Europe of accessions from both MA and A gene pools, but this is not observed in the comparisons between wilds vs domesticated populations.

#### GWAS on photoperiod sensitivity in the scarlet runner bean

Genome-wide association of flowering time in *Phaseolus coccineus* under long- and short-day conditions, using 666,489 sites (maf>0.05) identified several significant SNPs collocating with genes known to regulate flowering in other legumes, many of which have not been previously described in runner bean. On chromosome 1, a significant association (Chr01_18639957, p-value 1.9e-23; maf=0.41) explaining 11.6% of the phenotypic variance was detected near FT (FLOWERING LOCUS T; Phvul.001G097300, position:18602292-18604694), a central integrator of photoperiodic signals that promotes floral transition by activating downstream floral meristem identity genes.

On chromosome 5, a SNP (Chr05_66387, p-value 1.81e-17, maf=0.06) was located close to VRN1, a MADS-box transcription factor that facilitates floral induction by repressing flowering inhibitors and promoting the expression of floral activators, suggesting potential cross-talk between vernalization-like pathways and photoperiod responses in runner bean.

Chromosome 9 harbored associations (SNP Chr09_15987904, p-value 1.90e-35, maf=0.05; Chr09_30870720, p-value 1.20e-19, maf=0.31) near FPF1 (FLOWERING PROMOTING FACTOR 1) and AGL8-FUL (Agamous-like MADS-box protein AGL8). *FPF1* is an early flowering promoter that accelerates floral transition downstream of photoperiod sensing, while *AGL8-FUL* coordinates meristem identity and floral organ development. Their detection highlights previously unrecognized components of flowering regulation in *P. coccineus*.

Finally, on chromosome 10, a significant SNP (Chr10_39808334, p-value 8.27e-36, maf=0.39) was located near NST1 (NAC SECONDARY WALL THICKENING PROMOTING FACTOR 1). Explaining 22.3% of the phenotypic variance. Although primarily studied for its role in secondary cell wall biosynthesis, *NST1* may influence flowering indirectly through developmental progression, making it a novel candidate for flowering regulation in runner bean. Additional associations were detected across the genome; however, they could not be linked to annotated flowering or photoperiod genes.

In contrast, in *P. coccineus*, LD blocks around significant SNPs near FT, VRN1, FPF1, AGL8-FUL, and NST1 were shorter and less pronounced, consistent with the rapid LD decay typical of this outcrossing species. Photoperiod-sensitive accessions showed little to no LD at the same loci (Figure 4). These observations indicate that the GWAS signals are supported by local haplotype structure, but the extent of LD differs markedly between species due to their contrasting mating systems. We did not find overlaps between these GWAS signals and high iSAFE values or Fst outliers between photoperiod sensitive/insensitive cultivars (Suppl. figure 4).

**Figure 4.**
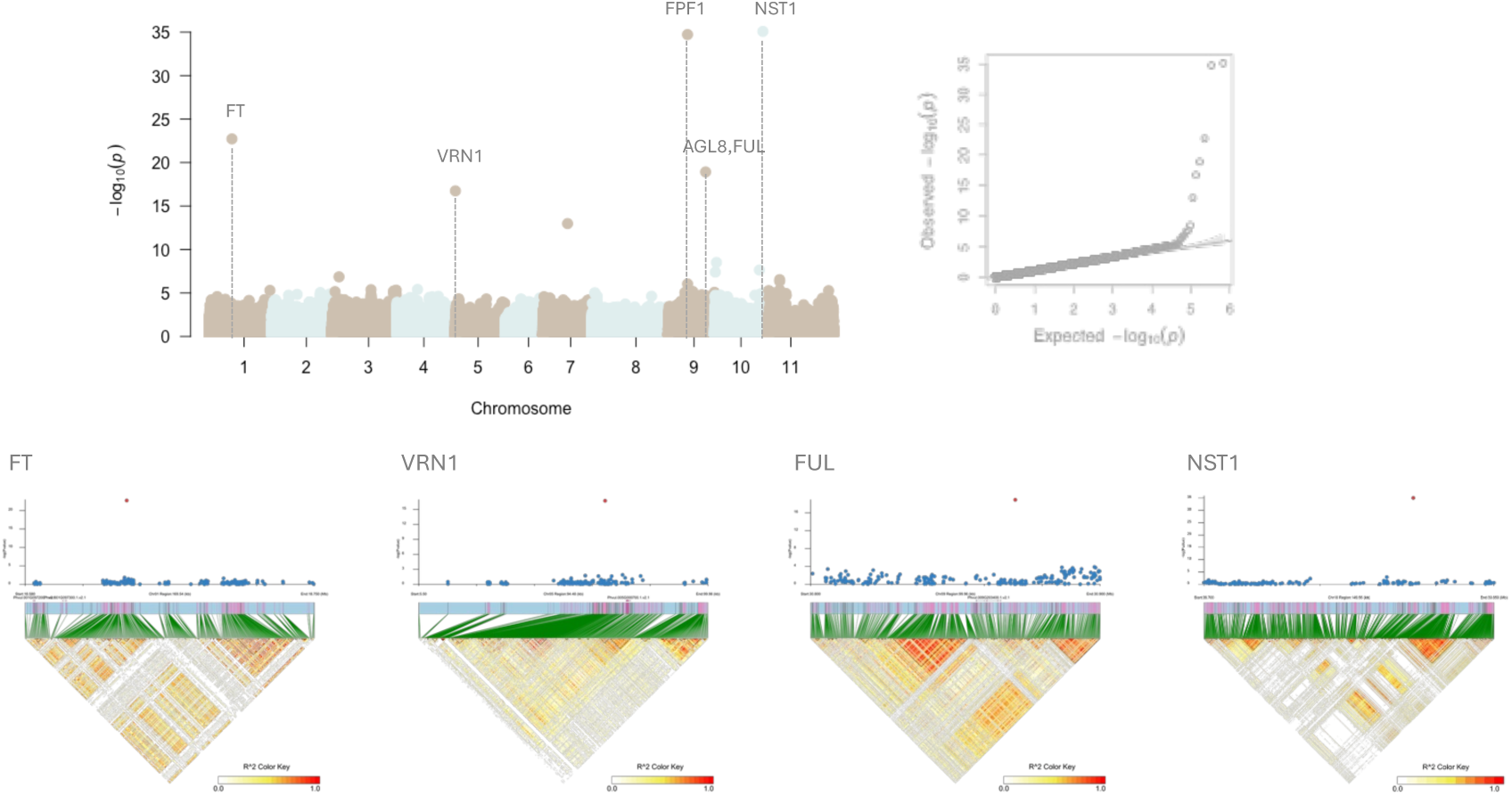
Genome-wide association analysis of photoperiod sensitivity in *Phaseolus coccineus*. (A) Manhattan plot showing SNP associations across the 11 chromosomes. GWAS was performed using the MLMM model implemented in GAPIT3. Significant peaks were detected on chromosomes 1, 5, 9, and 10. On chromosome 1, signals were located near FT, a central integrator of photoperiod cues; on chromosome 5, near VRN1, a MADS-box transcription factor involved in floral induction; on chromosome 9, near FPF1 and AGL8-FUL, which regulate meristem identity and floral organ development; and on chromosome 10, near NST1, a regulator of secondary cell wall formation potentially influencing flowering indirectly. (B) Quantile–quantile (Q–Q) plot of observed versus expected *p*-values, showing good model fit with enrichment of significant associations at the tail. (C) Linkage disequilibrium (LD) heatmaps and haplotype block structures for the main GWAS peaks. In contrast to *P. vulgaris*, *P. coccineus* exhibits short haplotype blocks around significant loci, consistent with its outcrossing mating system and rapid LD decay, which disperses associations across smaller genomic intervals. These patterns highlight the influence of reproductive biology on GWAS resolution and the genetic architecture of photoperiod adaptation. PhIns (left panel); PhSens (right panel).

## Discussion

The extent of parallel evolution during domestication and diversification has been extensively discussed but with no general agreement. Some authors argue there is little evidence for parallelism at the genetic level and that similar traits underlying the domestication syndrome in different species are usually controlled by loci that are not homologous (Glemin and Bataillon 2009; Martinez-Ainsworth and Tenaillon 2016). Other lines of evidence argue that pleiotropy has allowed for parallel evolution since most domestication genes appear to encode regulatory proteins affecting more than one trait (Sang 2009). Gaut (Gaut 2015) concluded that the question of the extent of parallel evolution during domestication of different crops remains open.

Flowering time in *Phaseolus vulgaris* and *P. coccineus* is governed by a complex interplay of photoperiod sensitivity and developmental determinacy. Our genome-wide association analyses revealed that, although both species have adapted to flower under the long days of Europe, they have done so by recruiting species-specific loci. In *P. vulgaris*, associations mapped near ELF4, E1, and LUX, while in *P. coccineus* we identified signals close to FT, VRN1, FPF1, and *AGL8-FUL*. These differences indicate that each species mobilized distinct genetic variants to meet the same ecological challenge.

Despite this divergence at the gene level, both bean species now display a similar adaptive phenotype: the capacity to flower under long-day conditions, a trait they began to express along domestication in the Americas and became nearly fixed after their introduction to Europe roughly 500 years ago. This highlights a case of rapid adaptation achieved through different allelic routes. Importantly, all associated loci belong to the photoperiod and circadian clock network, showing that while the exact genes differ, the evolutionary target is the same functional pathway. We interpret this as convergent evolution at the pathway level: different species-specific solutions that ultimately act on equivalent regulatory modules to ensure floral induction under long days. This kind of convergent trajectories has been discussed in other crops. For instance, the study by Larter et al. (2018) demonstrated that transitions in floral pigmentation across the Andean clade of *Iochrominae* involved predictable regulatory changes in the anthocyanin biosynthetic pathway, particularly in the expression of late pathway genes like F3’5’H, DFR, and ANS (Larter, et al. 2018). Similarly, our GWAS analyses identified consistent loci associated with photoperiod sensitivity and determinacy in both species, suggesting that despite their divergent mating systems, both species have undergone parallel regulatory adaptations to optimize reproductive timing. This convergence is likely constrained by the modular architecture of flowering pathways, where pleiotropic effects and regulatory interactions channel evolutionary changes along certain trajectories. Such pathway-level convergence highlights the predictability of evolutionary outcomes in response to similar selective pressures, even across species with different reproductive strategies.

Interestingly, the Ppd locus, long recognized as a central regulator of photoperiod sensitivity in beans (Weller, et al. 2019), did not produce any significant GWAS signals in our panel. However, our haplotype analysis revealed that linkage disequilibrium around *PHYA3* (the positional candidate for Ppd) is very high in photoperiod-insensitive cultivars (Suppl. Figure 5). This suggests that the lack of signal is not due to absence of association, but rather to the fact that the key adaptive alleles at Ppd are already nearly fixed in European germplasm. In other words, the selective sweep that allowed beans to adapt to long days may have already erased within-population variation at this locus, making it invisible to GWAS.

This pattern contrasts with findings in other crops, where convergence has sometimes been observed at the same gene or even nucleotide level. For example, in soybean, rice, and tomato, the *G* gene family has been repeatedly targeted during domestication to control seed dormancy (Wang, et al. 2018). Likewise, in lychee, independent domestication events converged on allelic variants of a single *Constans*-like gene that controls maturation timing (Hu, et al. 2022). In *Phaseolus*, however, we find no evidence for such gene-level convergence. Instead, adaptation to European long days has proceeded through species-specific gene recruitment, with convergence occurring at the broader functional scale of the photoperiod pathway. This underlines that parallel evolution in crop species can occur at different organizational levels, from single genes to entire regulatory networks.

A key methodological insight from our study concerns the Fin locus (TFL1 homolog). TFL1, a floral repressor and major regulator of determinacy, emerged as an overwhelmingly strong GWAS signal. If determinacy was not included as a co-variate, the association around TFL1 dominated our analyses and masked other important flowering loci. Only by explicitly modeling determinacy could we uncover additional genetic signals contributing to long-day adaptation. This demonstrates the dual role of TFL1: while central to growth habit variation, it also modulates flowering time, making it a dominant factor in the genetic architecture of phenology. Properly accounting for this pleiotropy was essential to disentangle the complexity of flowering responses in *Phaseolus*.

The divergent mating systems also help explain the distinct haplotype patterns observed around major GWAS signals. In *P. vulgaris*, the reduced recombination inherent to selfing maintains extended regions of LD, resulting in long haplotype blocks surrounding dominant signals. These extended haplotypes facilitate the detection of strong associations but also increase the likelihood of linked, non-causal variants contributing to the signals. By contrast, in *P. coccineus*, high outcrossing rates and rapid LD decay prevent the formation of long haplotype blocks, leading to more localized and fragmented association signals. This explains why no extended haplotype structures were detected around significant loci in *P. coccineus*, despite its clear adaptation to long-day flowering. Together, these patterns underscore how reproductive biology shapes the genomic architecture of adaptation and the resolution of GWAS analyses.

Taken together, our results show that *P. vulgaris* and *P. coccineus* have both undergone rapid adaptation to long-day flowering, but through species-specific genetic routes. The convergence resides not at the locus or allelic level, but at the pathway level, underscoring the centrality of the photoperiod regulatory network as the evolutionary solution to the constraints of temperate latitudes. This broader, pathway-scale convergence provides a compelling example of how different species can follow distinct molecular trajectories yet arrive at equivalent adaptive outcomes.

## Acknowledgements

The common bean collection was phenotyped at the Plant Cultivation Facility in Biocentrum, SLU campus Ultuna. We want to thank the team in charge of the phytotrons and green-house, Urban Pettersson, Per Linden, Fredric Hedlund and Kathrin Hesse. This research was financed by the Swedish Research Council (VR), under the grant no. 2018-03780 to PKI. We acknowledge support from the National Genomics Infrastructure in Stockholm funded by Science for Life Laboratory, the Knut and Alice Wallenberg Foundation and the Swedish Research Council for assistance with massively parallel sequencing. All computational analyses and data handling were enabled by resources provided by the Swedish National Infrastructure for Computing (SNIC) at Uppsala Multidisciplinary Centre for Advanced Computational Science (UPPMAX) under the computing projects 2019/3-597, 2020/5-621 and storage project sllstore2017050.

## Data accessibility

Raw read data have been uploaded to NCBI and can be found under the SRA Bioproject PRJNA1004188.

## Author contributions

MR-A planned and designed the research. PKI provided scientific feedback. MR-A, XF and LY performed experiments. MR-A wrote the manuscript. All authors read and approved of the final version of the manuscript.

## References

1. Akbari A, Vitti JJ, Iranmehr A, Bakhtiari M, Sabeti PC, Mirarab S, Bafna V. 2018. Identifying the favored mutation in a positive selective sweep. Nat Methods 15:279–282.

2. Angioi SA, Rau D, Attene G, Nanni L, Bellucci E, Logozzo G, Negri V, Spagnoletti Zeuli PL, Papa R. 2010a. Beans in Europe: origin and structure of the European landraces of Phaseolus vulgaris L. Theoretical and Applied Genetics 121:829–843.

3. Angioi SA, Rau D, Attene G, Nanni L, Bellucci E, Logozzo G, Negri V, Spagnoletti Zeuli PL, Papa R. 2010b. Beans in Europe: origin and structure of the European landraces of Phaseolus vulgaris L. Theor Appl Genet 121:829–843.

4. Blackman BK. 2017. Changing Responses to Changing Seasons: Natural Variation in the Plasticity of Flowering Time. Plant Physiol 173:16–26.

5. Browning SR, Browning BL. 2007. Rapid and accurate haplotype phasing and missing-data inference for whole-genome association studies by use of localized haplotype clustering. Am J Hum Genet 81:1084–1097.

6. Fang C, Sun Z, Li S, Su T, Wang L, Dong L, Li H, Li L, Kong L, Yang Z, et al. 2024. Subfunctionalisation and self-repression of duplicated E1 homologues finetunes soybean flowering and adaptation. Nat Commun 15:6184.

7. Gaut BS. 2015. Evolution Is an Experiment: Assessing Parallelism in Crop Domestication and Experimental Evolution: (Nei Lecture, SMBE 2014, Puerto Rico). Mol Biol Evol 32:1661–1671.

8. Glemin S, Bataillon T. 2009. A comparative view of the evolution of grasses under domestication. New Phytol 183:273–290.

9. Gonzalez AM, Vander Schoor JK, Fang C, Kong F, Wu J, Weller JL, Santalla M. 2021. Ancient relaxation of an obligate short-day requirement in common bean through loss of CONSTANS-like gene function. Curr Biol 31:1643–1652 e1642.

10. Gonzalez AM, Yuste-Lisbona FJ, Saburido S, Bretones S, De Ron AM, Lozano R, Santalla M. 2016. Major Contribution of Flowering Time and Vegetative Growth to Plant Production in Common Bean As Deduced from a Comparative Genetic Mapping. Front Plant Sci 7:1940.

11. Hu G, Feng J, Xiang X, Wang J, Salojarvi J, Liu C, Wu Z, Zhang J, Liang X, Jiang Z, et al. 2022. Two divergent haplotypes from a highly heterozygous lychee genome suggest independent domestication events for early and late-maturing cultivars. Nat Genet 54:73–83.

12. Koinange EMK, Singh SP, Gepts P. 1996. Genetic Control of the Domestication Syndrome in Common Bean. Crop Science 36:cropsci1996.0011183X003600040037x.

13. Korneliussen TS, Albrechtsen A, Nielsen R. 2014. ANGSD: Analysis of Next Generation Sequencing Data. BMC Bioinformatics 15:356.

14. Kwak M, Toro O, Debouck DG, Gepts P. 2012. Multiple origins of the determinate growth habit in domesticated common bean (Phaseolus vulgaris). Ann Bot 110:1573–1580.

15. Kwak M, Velasco D, Gepts P. 2008. Mapping homologous sequences for determinacy and photoperiod sensitivity in common bean (Phaseolus vulgaris). J Hered 99:283–291.

16. Larter M, Dunbar-Wallis A, Berardi AE, Smith SD. 2018. Convergent Evolution at the Pathway Level: Predictable Regulatory Changes during Flower Color Transitions. Mol Biol Evol 35:2159–2169.

17. Li H. 2011. A statistical framework for SNP calling, mutation discovery, association mapping and population genetical parameter estimation from sequencing data. Bioinformatics 27:2987–2993.

18. Li H, Durbin R. 2009. Fast and accurate short read alignment with Burrows-Wheeler transform. Bioinformatics 25:1754–1760.

19. Li H, Handsaker B, Wysoker A, Fennell T, Ruan J, Homer N, Marth G, Abecasis G, Durbin R, Genome Project Data Processing S. 2009. The Sequence Alignment/Map format and SAMtools. Bioinformatics 25:2078–2079.

20. Liew LC, Singh MB, Bhalla PL. 2017. A novel role of the soybean clock gene LUX ARRHYTHMO in male reproductive development. Sci Rep 7:10605.

21. Martinez-Ainsworth NE, Tenaillon MI. 2016. Superheroes and masterminds of plant domestication. C R Biol 339:268–273.

22. Pickersgill B. 2018. Parallel vs. Convergent Evolution in Domestication and Diversification of Crops in the Americas. Frontiers in Ecology and Evolution 6.

23. Purcell S, Neale B, Todd-Brown K, Thomas L, Ferreira MA, Bender D, Maller J, Sklar P, de Bakker PI, Daly MJ, et al. 2007. PLINK: a tool set for whole-genome association and population-based linkage analyses. Am J Hum Genet 81:559–575.

24. Rodriguez M, Rau D, Angioi SA, Bellucci E, Bitocchi E, Nanni L, Knupffer H, Negri V, Papa R, Attene G. 2013. European Phaseolus coccineus L. landraces: population structure and adaptation, as revealed by cpSSRs and phenotypic analyses. PLoS One 8:e57337.

25. Roux F, Touzet P, Cuguen J, Le Corre V. 2006. How to be early flowering: an evolutionary perspective. Trends in Plant Science 11:375–381.

26. Sang T. 2009. Genes and mutations underlying domestication transitions in grasses. Plant Physiol 149:63–70.

27. Schmutz J, McClean PE, Mamidi S, Wu GA, Cannon SB, Grimwood J, Jenkins J, Shu S, Song Q, Chavarro C, et al. 2014. A reference genome for common bean and genome-wide analysis of dual domestications. Nat Genet 46:707–713.

28. Skotte L, Korneliussen TS, Albrechtsen A. 2013. Estimating individual admixture proportions from next generation sequencing data. Genetics 195:693–702.

29. Wang F, Han T, Jeffrey Chen Z. 2024. Circadian and photoperiodic regulation of the vegetative to reproductive transition in plants. Commun Biol 7:579.

30. Wang J, Zhang Z. 2021. GAPIT Version 3: Boosting Power and Accuracy for Genomic Association and Prediction. Genomics Proteomics Bioinformatics 19:629–640.

31. Wang M, Li W, Fang C, Xu F, Liu Y, Wang Z, Yang R, Zhang M, Liu S, Lu S, et al. 2018. Parallel selection on a dormancy gene during domestication of crops from multiple families. Nat Genet 50:1435–1441.

32. Weller JL, Ortega R. 2015. Genetic control of flowering time in legumes. Front Plant Sci 6:207.

33. Weller JL, Vander Schoor JK, Perez-Wright EC, Hecht V, Gonzalez AM, Capel C, Yuste-Lisbona FJ, Lozano R, Santalla M. 2019. Parallel origins of photoperiod adaptation following dual domestications of common bean. J Exp Bot 70:1209–1219.

34. Woodhouse MR, Hufford MB. 2019. Parallelism and convergence in post-domestication adaptation in cereal grasses. Philos Trans R Soc Lond B Biol Sci 374:20180245.

35. Wu T, Liu Z, Yu T, Zhou R, Yang Q, Cao R, Nie F, Ma X, Bai Y, Song X. 2024. Flowering genes identification, network analysis, and database construction for 837 plants. Hortic Res 11:uhae013.

36. Zhang C, Dong SS, Xu JY, He WM, Yang TL. 2019. PopLDdecay: a fast and effective tool for linkage disequilibrium decay analysis based on variant call format files. Bioinformatics 35:1786–1788.

