## Supplementary material for "Species-specific genetic routes to a shared phenotype: convergent adaptation of beans to European long days": Suppl. Figure 1

**Supplementary figures.**

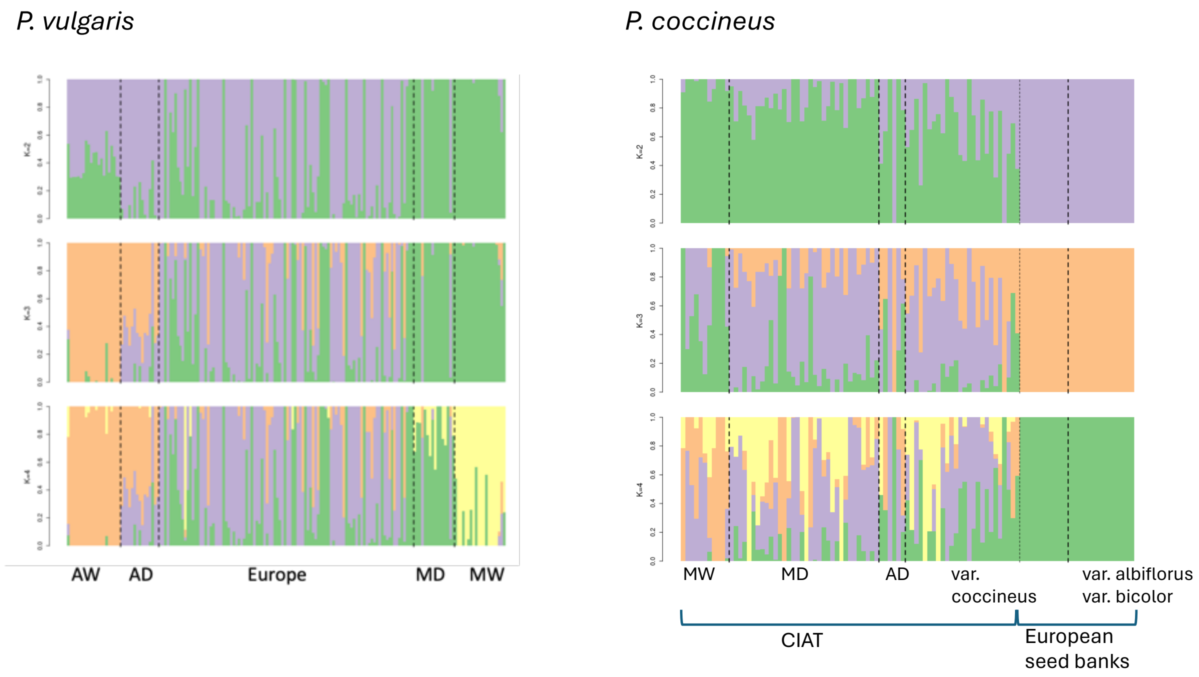


**Suppl. Figure 1.** NGSAdmix plots for the common and scarlet runner bean.

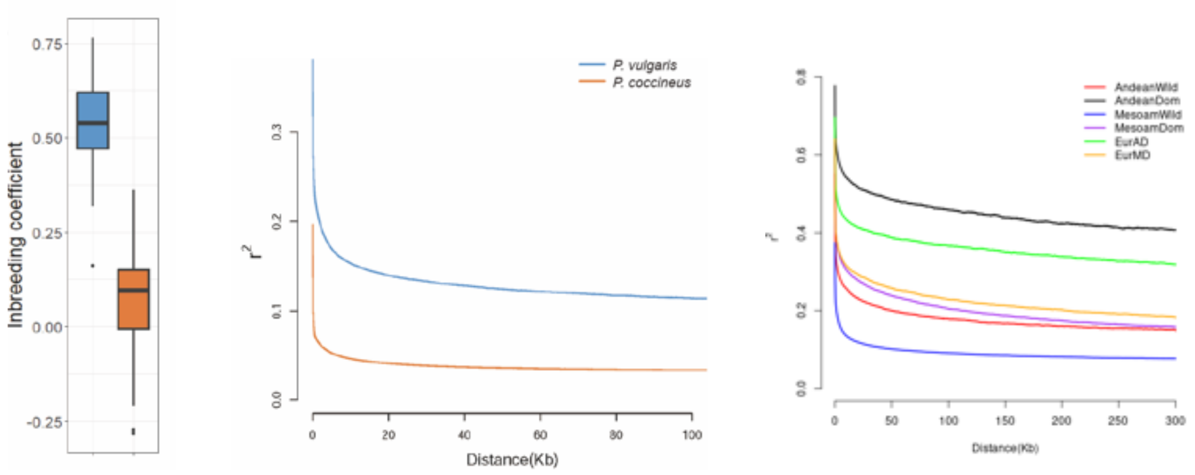


**Suppl. Figure 2.** Linkage disequilibrium decay and inbreeding coefficients in *Phaseolus vulgaris* and *P. coccineus*. (A) Distribution of inbreeding coefficients (F) across accessions of both species. *P. vulgaris* displays high F values, consistent with predominant selfing, while *P. coccineus* shows values near zero, consistent with extensive outcrossing. (B) Decay of linkage disequilibrium (LD) with physical distance between SNPs for both species. *P. vulgaris* (selfing) shows slow LD decay, with $r^2$ values remaining high over long distances, whereas *P. coccineus* (outcrossing) exhibits rapid LD decay, with $r^2$ dropping sharply within short physical distances. The contrasting patterns reflect the differing recombination histories and mating systems of the two species.

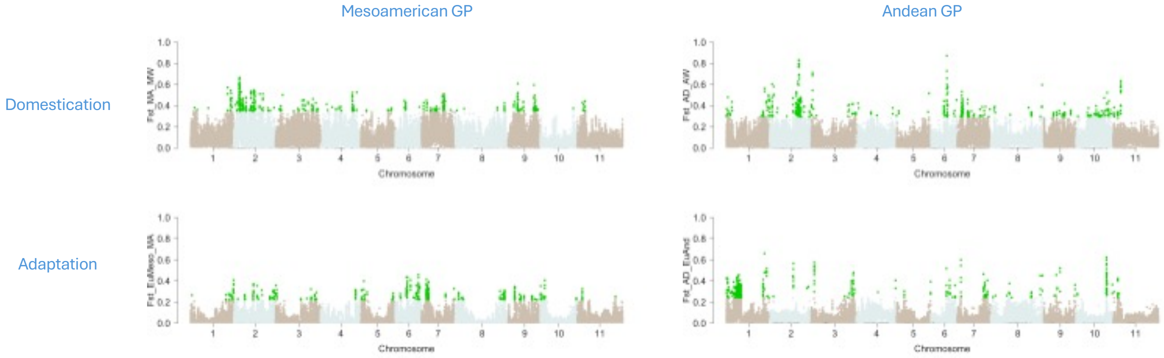


**Suppl. Figure 3.** Genome-wide differentiation between *Phaseolus vulgaris* populations. Manhattan plots displaying Fst values calculated between major population groups of *P. vulgaris*, in 10Kb, non-overlapping windows. Outliers (highlighted in green) correspond to the top 5% Fst values. The genome-wide distribution highlights both highly differentiated regions as a result of independent domestication and further adaptation to Europe.

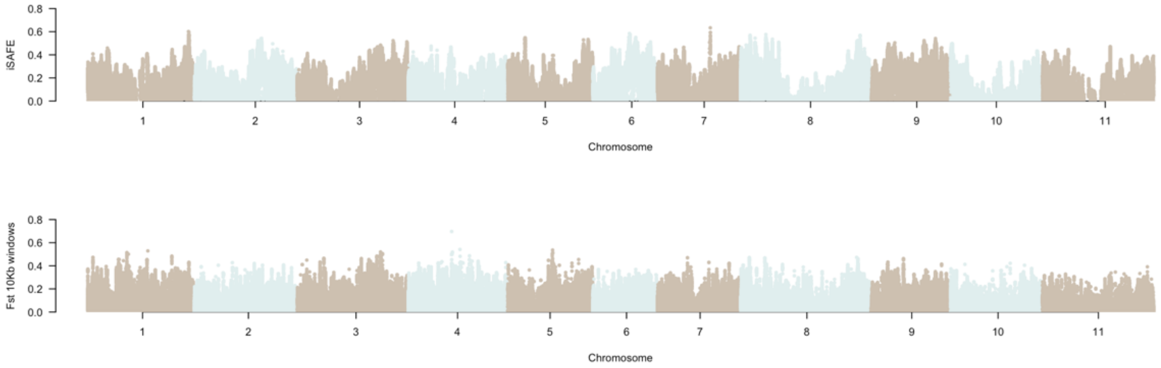


**Suppl. Figure 4.** Detection of selective sweeps and population differentiation in *Phaseolus coccineus*. Genome-wide signals of selection were evaluated using iSAFE (integrated Selection of Allele Favored by Evolution) and population differentiation Fst. (A) iSAFE values identify regions likely under positive selection in PhIns. (B) Fst values between PhIns/PhSens population groups.

Phytochrome A

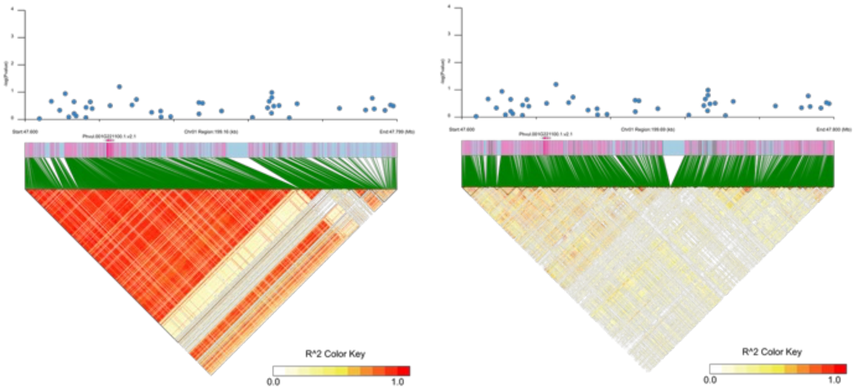


**Suppl. Figure 5.** Linkage disequilibrium (LD) heatmap and haplotype block structure around the PhyA locus in chromosome 1 in PhIns (left panel) and PhSens (right panel) *P. vulgaris*.
